# Activity Dependent Stimulation Increases Synaptic Efficacy In Spared Pathways In An Anesthetized Rat Model Of Spinal Cord Contusion Injury

**DOI:** 10.1101/2020.06.11.146910

**Authors:** Jordan A. Borrell, Dora Krizsan-Agbas, Randolph J. Nudo, Shawn B. Frost

**Affiliations:** Bioengineering Program, University of Kansas, Lawrence, KS, USA; Landon Center on Aging, University of Kansas Medical Center, Kansas City, KS, USA; Molecular & Integrative Physiology, University of Kansas Medical Center, Kansas City, KS, USA; Department of Rehabilitation Medicine, University of Kansas Medical Center, Kansas City, KS, USA

**Keywords:** Activity Dependent Stimulation, Neuromodulation, Spinal Cord Injury, Extracellular Electrophysiology, Synaptic Efficacy

## Abstract

The purpose of this study was to assess the ability of intraspinal microstimulation (ISMS), triggered by action potentials (spikes) recorded in motor cortex, to alter synaptic efficacy in descending motor pathways in an anesthetized rat model of spinal cord injury (SCI). Experiments were carried out in adult, male, Sprague Dawley rats with a moderate contusion injury at T8. For activity-dependent stimulation (ADS) sessions, a recording microelectrode was used to detect neuronal spikes in motor cortex that triggered ISMS in the spinal cord grey matter. SCI rats were randomly assigned to one of four experimental groups differing by: a) cortical spike-ISMS stimulus delay (10 or 25 ms) and b) number of ISMS pulses (1 or 3). Four weeks after SCI, ADS sessions were conducted in three consecutive 1-hour conditioning bouts for a total of 3 hours. At the end of each conditioning bout, changes in synaptic efficacy were assessed using intracortical microstimulation (ICMS) to examine the number of spikes evoked in spinal cord neurons during 5-minute test bouts. A multichannel microelectrode recording array was used to record cortically-evoked spike activity from multiple layers of the spinal cord. The results showed that ADS resulted in an increase in cortically-evoked spikes in spinal cord neurons at specific combinations of spike-ISMS delays and numbers of pulses. Efficacy in descending motor pathways was increased throughout all dorsoventral depths of the hindlimb spinal cord. These results show that after an SCI, ADS can increase synaptic efficacy in spared pathways between motor cortex and spinal cord. This study provides further support for ADS as an effective therapeutic approach for enhancing descending motor control after SCI.

## Introduction

Closed-loop neuromodulation systems have received increased attention in recent years as potential therapeutic approaches for treating neurological injury and disease. These systems utilize various signal sources, such as EMG, to trigger electrical stimulation of the spinal cord. For example, intraspinal microstimulation (ISMS), triggered by volitional muscle activity in the impaired forelimb can result in improved behavioral performance of the forelimb in a rat model of cervical spinal cord injury [1]. ISMS is synchronized with the arrival of motor commands signaled by EMG activity, presumably enhancing synaptic plasticity. In another variation of activity-dependent stimulation (ADS), EMG-triggered epidural electrical stimulation resulted in improvement in motor control of locomotion in rat models of complete spinal cord transection [2, 3].

ADS paradigms using recorded spikes from motor cortex to trigger spinal cord stimulation may allow for much greater temporal precision and thus, may result in more robust synaptic plasticity. Recently, it was demonstrated that epidural spinal cord stimulation triggered by neuronal spikes from the hindlimb motor cortex could restore weight-bearing locomotion of the paralyzed leg in non-human primates with a unilateral spinal cord transection [4]. However, the primary rationale for this study was not to enhance synaptic plasticity in the spinal cord, per se, but to provide direct control over hindlimb muscle activity. More precise control over the temporal coupling of motor command signals and depolarization of spinal cord motor neurons may provide a means for enhancing synaptic plasticity [5] and thus, for restoring motor function.

ADS paradigms specifically based on the timing of pre- and post-synaptic events are beginning to be tested in injury models to determine their potential clinical utility in strengthening remaining intact connections. Replication of the natural timing of pre- and post-synaptic events has been shown to potentiate, or condition, neuronal connections *in vivo,* resulting in functional plasticity of motor cortex outputs [6]. Spike-timing dependent plasticity has been shown to result in long-term changes in the strength of corticospinal connections in non-human primates that can last for at least a few days [7].

The purpose of the present study was to determine whether greater control over spike-stimulus delays can increase synaptic efficacy in remaining, intact descending motor pathways after a lower thoracic spinal cord contusion injury. Spike-timing dependence is a particularly important issue for SCI applications of ADS, since motor cortex outputs influence spinal cord neurons via various descending pathways (e.g., corticospinal, cortico-reticulospinal pathways) with different conduction velocities and multiple synaptic connections. Our hypothesis is that the synaptic efficacy of spared descending pathways from motor cortex to the spinal cord can be strengthened by using ADS to replicate the natural timing of pre- and post-synaptic events, resulting in an increase in cortically-evoked activity in the hindlimb spinal cord. These initial studies were performed in an acute anesthetized rat model four weeks after a lower thoracic contusion injury. We used spike activity recorded in motor cortex to precisely time ISMS relative to motor cortex outputs. The effect of ADS on synaptic efficacy below the level of the contusion injury was measured by changes in neuronal spikes recorded in the spinal cord that were evoked by intracortical microstimulation. The anesthetized preparation allowed us to record neuronal activity throughout the depths of the spinal cord using linear multielectrode arrays.

The results demonstrate that cortically driven ISMS strengthens the synaptic efficacy of spared descending pathways after lower thoracic contusion injury, and that different populations of spinal cord neurons are differentially strengthened by different time delays. This application of ADS (i.e., cortical spike-driven ISMS) in an animal model of thoracic spinal cord contusion injury has not been previously studied. The results provide further support for synaptic plasticity induced by the therapeutic use of precisely-timed ADS after SCI.

## Methods

### Subjects

Thirty-four adult, male, Sprague Dawley rats were used in this study. Body weights ranged from 356—486 g (411.1 ± 29.8 g) and ages ranged from 90—105 days old (96.0 ± 4.0 days old) at the time of the electrophysiological experiments. Each rat underwent a surgical procedure to induce a midline spinal cord contusion injury at T8. Four weeks later, rats underwent neurophysiological procedures to examine the effects of two ADS stimulation parameters: 1) The time delay between recorded action potentials (i.e., spikes) in the motor cortex hindlimb area (HLA) and onset of intraspinal microstimulation (ISMS) in the ventral horn of the lumbar spinal cord (10 or 25 ms) and 2) the number of biphasic ISMS pulses (1 or 3). Ten rats were removed from the study due to adverse surgical complications. Thus, 24 rats were randomly assigned to one of four ADS parameter groups and completed the study: 1) *Group 10ms_1P* (n = 7): 10 ms time delay with 1 ISMS pulse; 2) *Group 10ms_3P* (n= 5): 10 ms time delay with 3 pulses; *Group 25ms_1P* (n = 7) 25 ms time delay with 1 pulse; and *Group 25ms_3P* (n = 5): 25 ms time delay with 3 pulses. All procedures were performed in accordance with the *Guide for the Care and Use of Laboratory Animals Eighth Edition* (Institute for Laboratory Animal Research, National Research Council, Washington, DC: National Academy Press, 2011). The protocol was approved by the University of Kansas Medical Center Institutional Animal Care and Use Committee.

### General Surgical Procedures

In each rat, two surgical procedures were performed: 1) a recovery surgical procedure to induce a contusion SCI and 2) a terminal procedure to a) implant microelectrodes into hindlimb motor cortex and spinal cord, b) implant EMG electrodes into selected hindlimb muscles, and c) conduct ADS neurophysiological sessions (Figure 1). In both surgical procedures, after a stable anesthetic state was reached using isoflurane anesthesia, ketamine hydrochloride (100 mg kg^-1^; IP)/xylazine (5 mg kg^-1^; IP) was delivered. Anesthesia was maintained with subsequent doses of ketamine (0.1 mL; IP or IM) and monitored via pinch and corneal responses. Additional doses of ketamine were administered if the rat reacted to a pinch of the forepaw/hindpaw or blinked after lightly touching the cornea. Aseptic conditions were maintained for both procedures.

**Figure 1.**
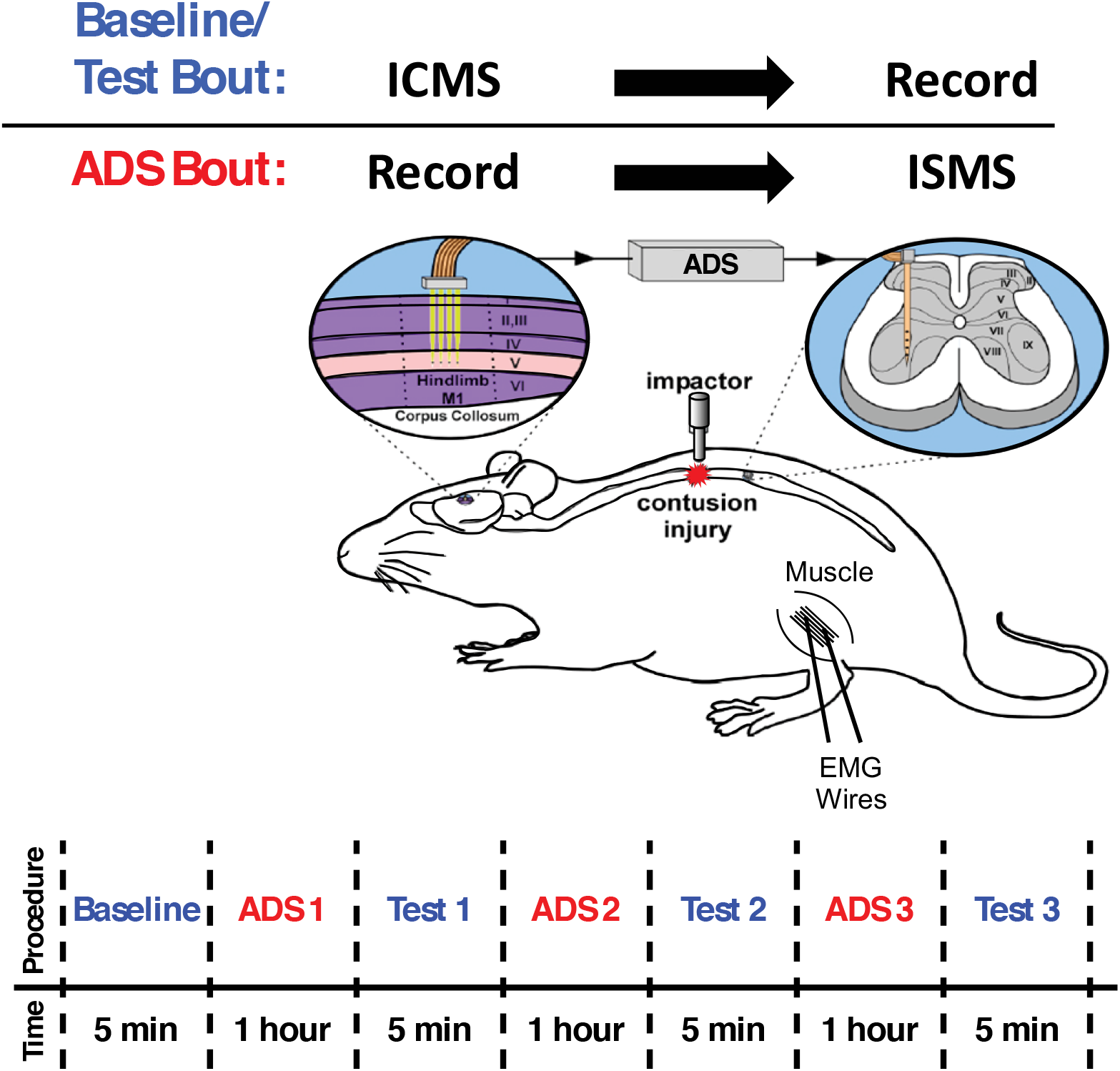
Overview of Experimental Design. Baseline/Test Bout: ICMS was applied to hindlimb motor cortex (HLA) while evoked spikes were recorded from spinal cord neurons below the level of a T8 contusion injury. **ADS Bout:** Spikes from HLA were recorded and used to trigger ISMS in the ventral horn of the thoracic spinal cord below the level of the injury. **ADS Timeline:** ADS bouts were conducted in one-hour sessions for a total of three hours. Test bouts were 5 min in duration, conducted at baseline and after each hour of ADS.

### Spinal cord injury (SCI)

All animals underwent a T8 laminectomy and moderate midline contusion injury using an Infinite Horizon spinal cord impactor (Precision Systems and Instrumentation, LLC, Fairfax Station, VA) with an impactor tip diameter of 2.5mm and 175 kDyn impact. Displacement distance reported by the impactor software for each contusion was recorded at the time of surgery and was used as an initial quantitative marker for successful impact. The average displacement value of the impactor was 1000.4 ± 123.8 μm (mean ± SD) from the surface of the spinal cord. At the conclusion of the surgery, 0.25% bupivacaine HCl was applied locally to the incision site. Buprenex (0.01 mg kg^-1^; SC) was administered immediately after surgery and 1 day later. All animals were monitored daily until the end of the experiment. On the day of surgery, and for 1-week post-surgery, the rats received a daily SC injection of 30,000 U of penicillin (Combi-Pen 48) in 5 mL of saline to prevent infections and dehydration. Rats’ bladders were expressed twice-daily until animals recovered urinary function. From the second week onward, animals were supplemented with vitamin C pellets (BioServ, Frenchtown, NJ) to avert urinary tract infection [8].

### BBB Scoring

Locomotor performance was assessed using the Basso, Beattie, and Bresnahan (BBB) scale [9]. BBB scoring was performed before SCI and once per week for up to 4 weeks post-SCI. Two investigators were always present for BBB scoring as recommended by Basso, et al. Four weeks was chosen because BBB scores typically plateau by 4 weeks and behavior is relatively stable [10, 11]. A straight walkway was utilized for the testing. Rats were habituated to the apparatus for 1 week prior to BBB scoring. A BBB score of 13-15 at 4 weeks post-SCI was set as the inclusion criterion for this study; however, no rats were removed from this study due to this criterion.

### Neurophysiology procedures

Experimental design. Neurophysiological sessions utilizing timing parameters based on mechanisms underlying spike-timing-dependent plasticity (STDP) were conducted four weeks after SCI. Spike-stimulus conditioning (i.e., ADS) was conducted in three, 1-hour ADS conditioning bouts for a total of 3 hours of ADS. Changes in synaptic efficacy were inferred by comparing spikes evoked by intracortical microstimulation (ICMS) during a 5 min baseline period before ADS and three 5 min test periods following each ADS bout (Figure 1).

#### EMG fine-wire electrode implantation

The implantation and verification of fine-wire EMG electrodes into hindlimb muscles was conducted as previously described [12]. EMG electrodes were implanted in four hindlimb muscles: the lateral gastrocnemius (LG), tibialis anterior (TG), vastus lateralis (VL), and biceps femoris (BF). Each EMG electrode consisted of a pair of insulated multi-stranded stainless-steel wires exposed approximately 1 mm, with the exposed, implanted end of the wire folded back on itself (‘hook’ electrode). Implantation locations were determined by surface palpation of the skin and underlying musculature. After the hindlimbs were shaved, EMG wires were inserted into the belly of each muscle with the aid of a 22-gauge hypodermic needle. For each EMG electrode pair, wires were positioned approximately 5 mm apart in each muscle. An additional ground lead was placed into the base of the tail. The external portion of the wires was secured to the skin with surgical glue (3M Vetbond Tissue Adhesive, St. Paul, MN) and adhesive tape.

To verify that the EMG electrodes were within the belly of the muscle, a stimulus isolator (BAK Electronics, Inc., Umatilla, FL) was used to deliver a biphasic (cathodic-leading) square-wave current pulse to the muscle through the implanted EMG electrodes. The impedance between the EMG electrodes was tested via an electrode impedance tester (BAK Electronics, Inc., Umatilla, FL). The EMG electrodes were determined to be inserted properly and within the desired muscle if the electrode impedance was 7-8 kΩ, the direct current delivery to the muscle resulted in contraction of the desired muscle and the movement threshold was ≤ 5mA.

#### Identification of motor cortex HLA

The rats were placed in a Kopf small-animal stereotaxic frame (David Kopf Instruments^®^, Tujunga, CA) and the incisor bar was adjusted until the heights of lambda and bregma were equal (flat skull position). The cisterna magna was punctured at the base of the skull to reduce edema during mapping. A craniectomy was performed over the motor cortex. The general location of the craniectomy was guided by previous motor mapping studies in the rat [13, 14]. The dura over the cranial opening was incised and the opening was filled with warm, medical grade, sterile, silicone oil (50% Medical Silicone Fluid 12,500, 50% MDM Silicone Fluid 1000, Applied Silicone Corp., Santa Paula, CA) to prevent desiccation during the experiment.

A magnified digital color photograph of the exposed cortex was taken through a surgical microscope. The image was transferred to Canvas 3.5 where a 200 μm grid was superimposed, by calibration with a millimeter ruler, to indicate intended sites for microelectrode penetration. ICMS was conducted in the HLA of the primary motor cortex (M1) to verify the topographic location (Figure 2A). In rats with contusion injuries as applied here, hindlimb movements cannot be elicited with ICMS from HLA. Thus, an indirect approach was taken to assure that the cortical microelectrodes were located in HLA. First, the general location of the HLA can be predicted based on stereotaxic coordinates from previous ICMS studies in healthy rats [13]. Further, hip movements are consistently evoked immediately caudal to the forelimb area (blue dots in Figure 2A) and lateral to the trunk area (light blue dots in Figure 2A). In rats with thoracic contusion injuries, ICMS can evoke forelimb and trunk movements [14].

**Figure 2.**
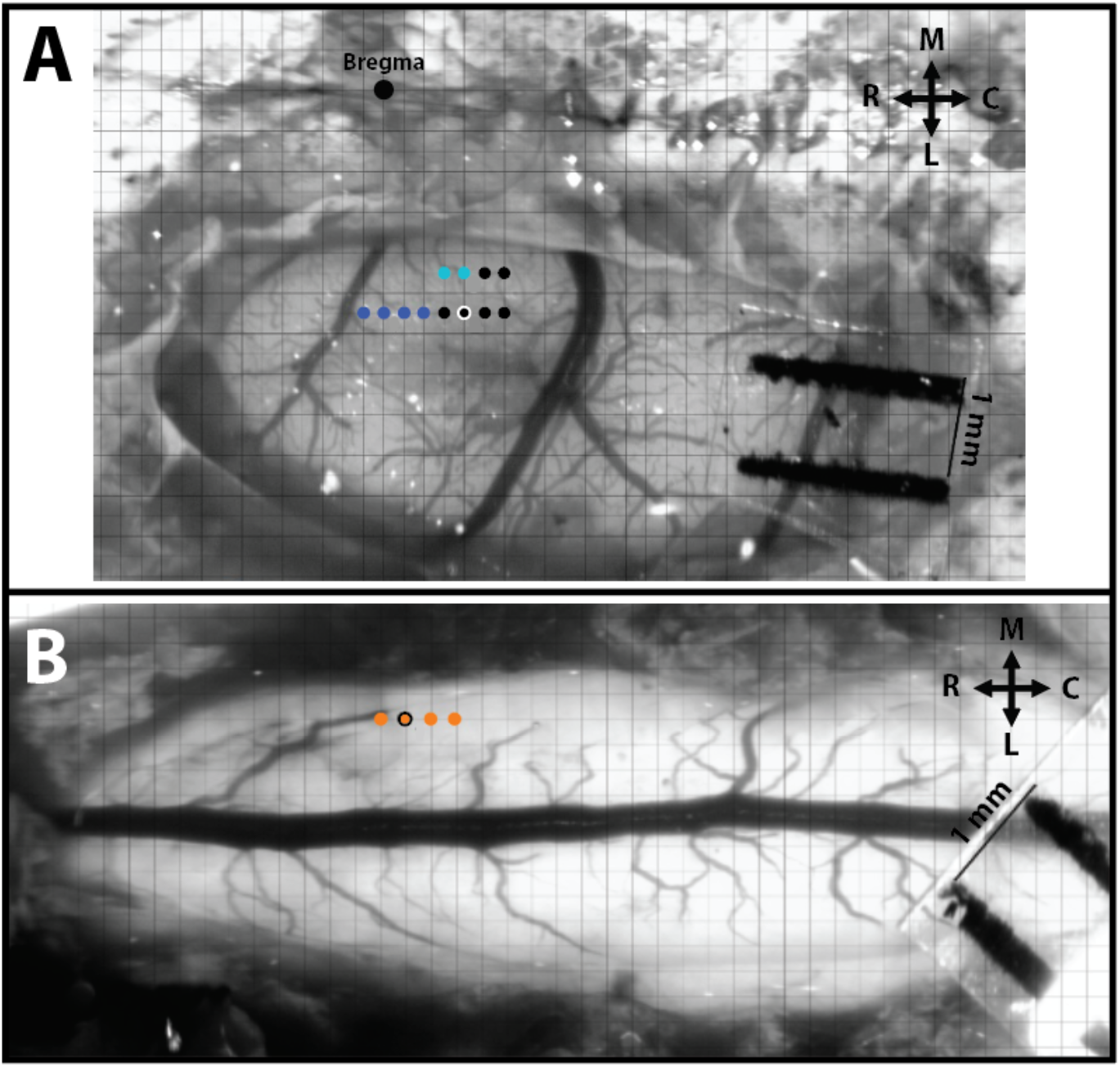
Intracortical and intraspinal microstimulation mapping and site pairing conducted in SCI rats. **A)** Hindlimb motor cortex with superimposed grid for ICMS mapping. The large black dot is at Bregma, the dark blue dots represent stimulation sites that resulted in ICMS-evoked forelimb movements, the light blue dots represent evoked trunk movements, and the small black dots represent sites of no evoked movement. **B)** Hindlimb spinal cord with superimposed grid for ISMS mapping. Each orange dot represents a stimulation site that resulted in ISMS-evoked hindlimb movement. The orange dot with the black outline represents the site of an ISMS-evoked hip movement paired for ADS. R = Rostral, C = Caudal, M = Medial, and L = Lateral.

ICMS was delivered to the ventral-most site on a single-shank Neuronexus probe (Neuronexus, Ann Arbor, MI) positioned at regularly spaced locations on the superimposed 200 μm grid. Each electrode site used for stimulation was iridium-activated and had an impedance in the range of ~50-70 kΩ with a surface area of 1250 μm^2^. The maximum compliance voltage of the system was 24 V. Electrode depth was controlled using a Kopf hydraulic microdrive (Kopf Instruments, Tujunga, CA) and reached ~1700 μm (i.e., Layer V of the cortex). The stimuli consisted of thirteen, 200 μs biphasic cathodal pulses delivered at 300 Hz repeated at 1/sec from an electrically isolated, charge-balanced (capacitively coupled) stimulation circuit. A maximum current of 100 μA was used during ICMS mapping.

#### Identification of lumbar spinal cord

Following ICMS mapping, a midline incision was made, exposing the T12-L2 vertebrate. A laminectomy was performed on the T13-L1 vertebrae exposing the L2-S1 segments of the spinal cord. The dura mater was removed using fine forceps and small scissors to allow electrode penetration. Animals were stabilized with a custom rodent spinal fixation frame (Keck Center for Collaborative Neuroscience Rutgers, The State University of New Jersey, USA) attached to the dorsal processes of vertebrae T12 and L2.

Previously derived 3-dimensional ISMS-evoked topographic maps were used in the guidance of stimulation sites in the hindlimb spinal cord [12]. A similar 200 μm grid was placed over a digital photomicrograph of the spinal opening for the mapping of evoked movements (Figure 2B). Stimulation sites were located ~0.8 mm lateral to the central blood vessel (i.e., midline) on the right side of the spinal cord and at a depth ~2.27 mm below the surface of the spinal cord (i.e., in the ventral horn). ISMS stimuli consisted of three 200 μs biphasic cathodal pulses delivered at 300 Hz repeated at 1/sec. ISMS-evoked movement threshold was measured during ISMS mapping and recorded as the lowest current level (i.e., increments of 1 μA) needed to evoke a consistent, repetitive joint displacement.

#### Intracortical and intraspinal evoked movement site matching

After ICMS was used to define forelimb and trunk sites, as well as non-responsive sites, a putative hip site (black dot with white outline in Figure 2A) was located. For subsequent ADS and test bouts, this site was paired with an ISMS-evoked hip site in the lumbar spinal cord (orange dot with black outline in Figure 2B).

#### ICMS-evoked spike recordings (Baseline and Test Bouts)

Using the paired sites of the HLA and hindlimb spinal cord, a stimulating electrode was inserted into the HLA and a recording microelectrode was simultaneously inserted in the hindlimb spinal cord site. The recording microelectrode was a single-shank, 16-channel Michigan-style linear microelectrode (Neuronexus, Ann Arbor, MI). Each of the 16 sites had a site area of 703 μm^2^, separation of 150 μm, thickness of 15 μm in diameter, and impedance of ~0.3 MΩ. The tip of the electrode was lowered to a depth of ~2270 μm below the spinal cord surface. ICMS was used to test corticospinal coupling by delivering a test stimulus in HLA, and ICMS-evoked spikes were recorded from the spinal cord and EMG from the hindlimb muscles. For baseline and test bouts, the ICMS stimulus consisted of three, 200 μs biphasic cathodal pulses delivered at 300 Hz repeated at 1/sec, and an ICMS intensity of 100 μA was used for all baseline and test bouts. During each baseline and test bout, neural and EMG activity were recorded and digitized for ~5 min from each of the 16 active recording sites and four fine-wire pairs, respectively, using neurophysiological recording and analysis equipment (Tucker Davis Technologies, Alachua, FL).

#### Activity-dependent-stimulation (ADS) paradigm

Using the paired HLA and spinal cord hip sites, the same recording microelectrode was inserted into the HLA site and the same stimulating microelectrode was inserted into the hindlimb spinal cord site. The stimulating microelectrode was positioned at ~2.27 mm below the surface of the spinal cord (i.e., in lamina VIII/IX in the ventral horn), as the long-range therapeutic goal is to enhance cortical modulation of spinal motor neurons. It is assumed that the stimulating electrode was either near spinal cord motor neuron cell bodies, the dendrites of motor neurons, or interneurons.

Using principal component analysis [15], spikes were recorded and sorted in real-time from a single channel in HLA (i.e., ~1700 μm below the surface in Layer V of the cortex) and used to trigger stimulation in the paired site in the hindlimb spinal cord. The triggered ISMS stimulus consisted of one or three, 200 μs biphasic cathodal pulses delivered at 300 Hz repeated at 1/sec. The stimulus intensity for ADS was set at 50% of the ISMS movement threshold in each rat to reduce the possibility of evoked movements during ADS therapy and to eliminate the possibility of fatigue on the muscle.

The ADS parameters of interest were time delay between recorded cortical spike and spinal cord stimulation and the number of ISMS pulses. Delays of 10 and 25 ms between recorded HLA spike and triggered stimulus pulse(s) were chosen based on natural response times observed in previous experiments [16]. Briefly, ICMS-evoked spiking activity in the hindlimb spinal cord was consistently observed with a 10 ms latency in healthy rats, hence a 10 ms time delay was chosen. A 25 ms time delay was chosen because it matched the latency of ICMS-evoked hindlimb EMG activity in healthy rats. Both time delays are known to fall within the window of long term potentiation [17] and fall within the window for altered cortical output responses when spike-triggered stimulation was used in motor cortex of non-human primates [6].

Three-pulse trains are typically used to evoke EMG activity and joint movements elicited by ISMS [18]. However, to document ISMS-evoked EMG activity more precisely, and to minimize polysynaptic effects, ADS effects were also tested using only one pulse as the ISMS stimulus.

### Neurophysiological data analysis

#### Spike sorting during baseline and test bouts

Neural spikes were discriminated offline with OpenSorter software (Tucker Davis Technologies, Alachua, FL) using principal component analysis [15]. ICMS-evoked activity recorded during the baseline and test bouts from each channel in the hindlimb spinal cord was represented in post-stimulus spike histograms for 200 ms around the onset of each ICMS stimuli (ICMS begins at 0 ms; histogram extends −100 ms before and +100 ms after onset ICMS) using custom software (Matlab; The Mathworks, Inc., Natick, MA). Spiking activity was averaged into 1 ms bins over the ~5 min recording (i.e., averaged over ~300 ICMS trains). Spike activity could not be discriminated during the first 7 ms after the onset of the ICMS train due to the stimulus artifact.

For analysis of spiking responses during the baseline and test bouts, the 16 recording channels in the hindlimb spinal cord were divided into four dorsoventral sectors based on depth below the surface of the spinal cord. The dorsoventral sectors (Figure 3) were: Dorsal Sector (720—1020 μm; dorsal horn), Upper Intermediate Sector (1120—1420 μm; intermediate layer), Lower Intermediate Sector (1520—1820 μm; intermediate layer), and Ventral Sector (1920—2220 μm; ventral horn).

**Figure 3.**
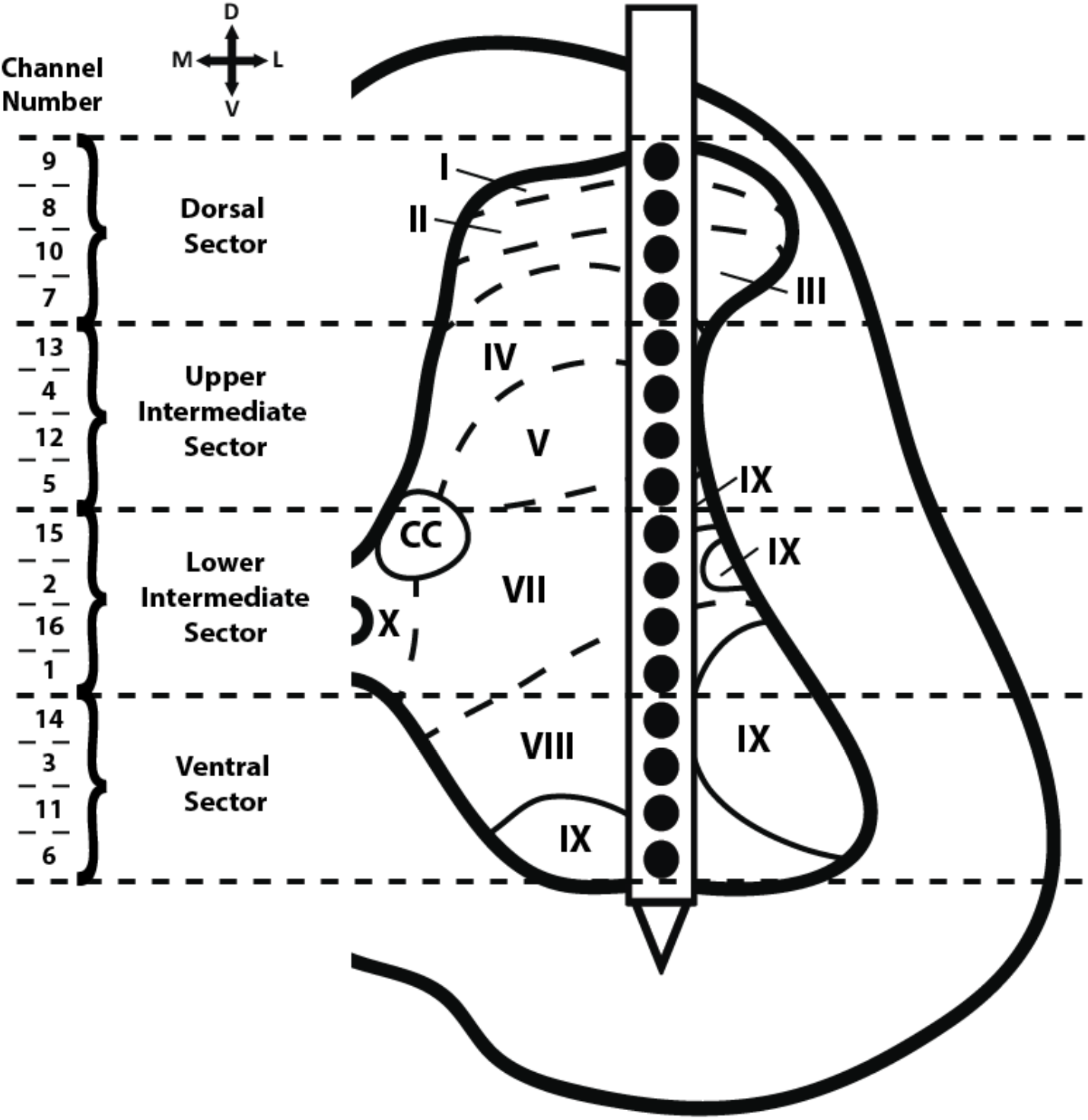
Cross-sectional schematic diagram of hindlimb spinal cord with placement of the recording electrode during the baseline and test bouts. The channel numbers of the recording sites in each of the four dorso-ventral sectors are shown at the left. Previous topographical studies indicate the rostral sector of vertebrae T13 of the Sprague Dawley rat is located at spinal cord segmental level L2-L3 [19]. Thus, the recording electrode was located in segmental level L2-L3. Based on cytoarchitectonic organization in the rat, electrode sites were in or near all laminae except for laminae VI (seen only in L3-S1 segments of the rat) and laminae X (found medially around the central canal) [20].

#### Firing rate (FR) calculation

The ICMS-evoked firing rate (FR) (i.e., total number of recorded spikes per 1ms bin/total time of recording) was calculated for each time bin of the post-stimulus spike histograms using custom software (Matlab; The Mathworks, Inc., Natick, MA). The FR was only considered for analysis if the FR for the respective time bin was greater than 2x standard deviation (horizontal, dashed, red line of Figure 7) above the average FR pre-ICMS (horizontal, solid, red line of Figure 7). Pre-ICMS FR was derived by averaging the FR of each time bin 10 ms before the onset of ICMS (i.e., from −10 to 0 ms in the post-stimulus spike histogram).

To narrow the range of potential cortical spike-ISMS delays, we utilized data from a previous study wherein we determined the latencies of ICMS-evoked action potentials in lower thoracic spinal cord neurons in healthy and SCI rats [16]. In healthy rats, ICMS-evoked spikes in the lower thoracic spinal cord occurred during a narrow, short-latency period (10-12 ms) as well as at longer latencies that formed a broader distribution (~20-40 ms). ICMS-evoked EMG activity was observed at ~25-28 ms post-ICMS. In SCI rats, ICMS-evoked spikes below the level of the SCI were found consistently at short latency (~10-12 ms). However, at the longer latencies, FR was substantially lower after the SCI, and only a few bins displayed FR significantly above baseline. Based on these previous results, the maximum FR was defined for the short-latency period (10-12 ms post-onset ICMS) and the long-latency period (24-36 ms). Then, the mean FR and standard error for the short- and long-latency periods were calculated for each baseline/test bout in each rat, and in each dorsoventral sector.

#### Stimulus-triggered average (StTA) of EMG

Using custom software (Matlab; The Mathworks, Inc., Natick, MA), StTAs of rectified EMG from leg muscles were plotted to determine muscle activation [12, 21]. For each of the implanted muscles, EMG data were recorded for at least 10 stimulus trains during ICMS and ISMS procedures at movement threshold or a maximum current of 100 μA and averaged over a time window of 220 ms. StTAs were aligned to the time of the first stimulus (i.e., 0 ms) and included data from −20.2 to +199.9 ms relative to the time of the first stimulus. A muscle was considered active when the average rectified EMG reached a peak ≥ 2.25 SD above baseline values in the interval from – 20.2 to 0 ms and had a total duration of ≥ 3 ms. The stimulus artifact was minimal to absent in EMG recordings with no muscle activation. If an artifact was observed, the amplitude of the artifact was minimal compared to the amplitude of the evoked EMG; however, the averaging of the StTAs of EMG recordings largely eliminated the stimulus artifact. As a result, the stimulus artifact was determined to have minimal to no effect on the recordings. EMG potentials were high- and low-pass filtered (30 Hz-2.5 kHz), amplified 200-1000 fold, digitized at 5 kHz, rectified and recorded on an RX-8 multi-channel processor (Tucker-Davis Technology, Alachua, FL).

### Statistical analyses

Statistical analyses were performed using JMP 11 software (SAS Institute Inc., Cary, NC). The nonparametric, Mann-Whitney U test was used to compare ordinal-scaled BBB scores. Bonferroni correction for multiple comparisons was used to control for multiple comparisons. ICMS-evoked spiking activity analyses followed the rationale of a comparable study [22]. The current study was designed to assess the effect and interactions of various ADS parameters in separate dorsoventral sectors over time on neuronal activity. Since the independence of individual neuron firing rates cannot be assumed within a particular rat and because the distribution of firing rates tends to have a Poisson distribution with a long tail (and thus, not normally distributed), a generalized linear mixed model (GLMM) for repeated measures was used. GLMM is a powerful statistical tool that permits modeling of not only the means of the data (as in the standard linear model) but also their variances as well as within-subject covariances (i.e., the model allows subjects with missing outcomes—unbalanced data—to be included in the analysis). Thus, data are presented as average ± standard error of the mean unless otherwise stated, and *p*-values < 0.05 were considered significant.

### Histology

At the end of each experiment, animals were euthanized with an intraperitoneal injection of sodium pentobarbital (Beuthanasia-D; 100 mg kg^-1^) and transcardially perfused with 4% paraformaldehyde in 0.1 M PBS. The spinal cord was sectioned at 30 μm on a cryostat in the coronal plane. Histology was performed on selected sections with cresyl violet stain for Nissl bodies to verify the injury.

## Results

### BBB Scores

Each rat scored 21 on the BBB scale (normal) before SCI. At 1-week post-SCI, BBB scores were significantly lower relative to baseline (Table I; *p* < 0.0001). BBB scores improved over subsequent weeks but remained significantly lower relative to baseline at 4 weeks post-SCI (Table 1; *p* < 0.0001). There was no statistical difference in BBB scores between groups for any week (*p* > 0.05). At 4-weeks post-SCI, BBB scores confirmed that rats had consistent weight-supported plantar steps, no or occasional toe clearance (i.e., scraping of the digits on the ground) during forward limb advancement, consistent forelimb-hindlimb coordination, and predominant paw position during locomotion that was always rotated externally just before it lifted off at the end of stance and either externally rotated or parallel to the body at initial contact with the surface.

**Table I.**
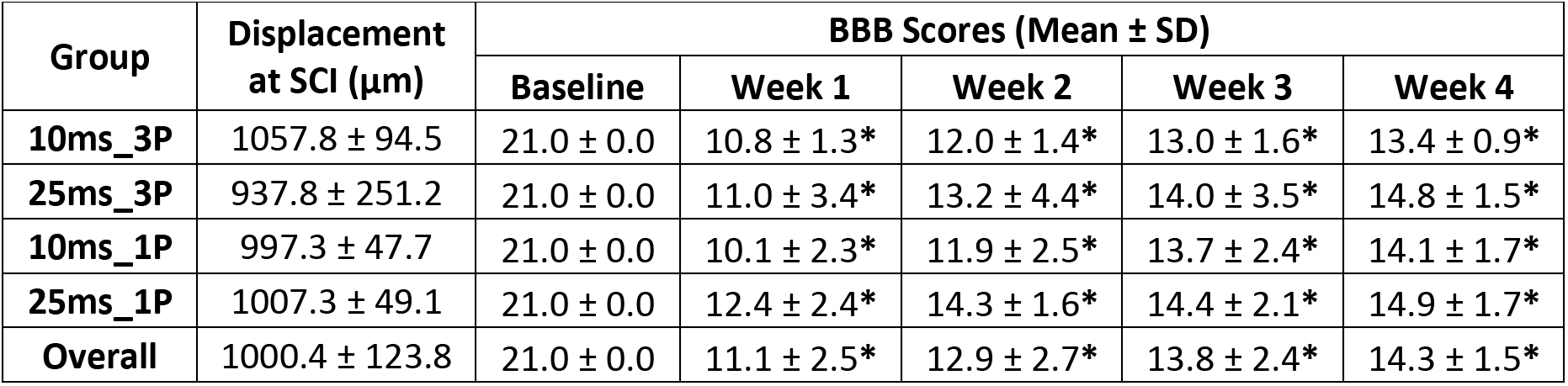
Behavioral Scores. Mean BBB score at baseline and at weeks 1-4 post-SCI and displacement of impactor at time of SCI for each rat group and all rats combined (+/- S.D.). There was no statistical difference in impactor displacement at SCI nor in BBB scores between groups at any of the four weeks (*p* > 0.05). * = Significant difference of BBB score when compared to baseline (within-group comparison; *p* < 0.0001).

### SCI Verification

A representative image of a histological section through the injury epicenter is shown in Figure 4. Briefly, the injury spared an outer rim of white matter that surrounded a central core lesion. Remnants of the gray matter were rarely seen at the lesion center.

**Figure 4.**
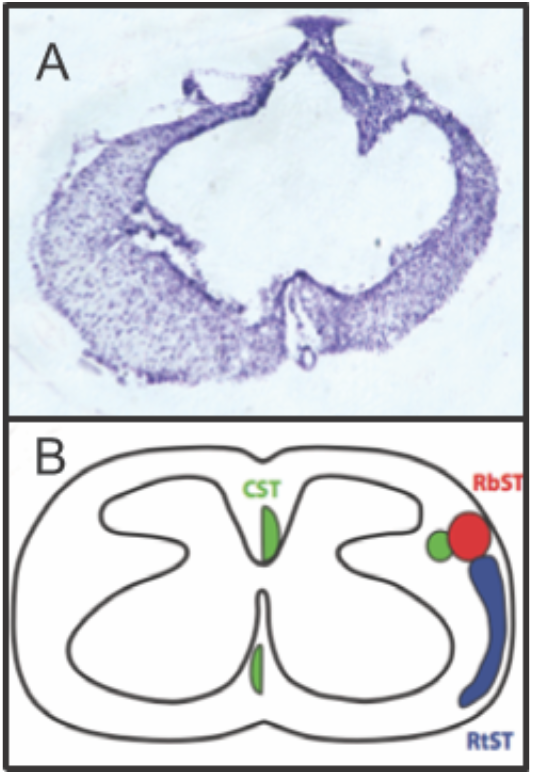
Representative image of the spinal cord injury epicenter 4 weeks after injury. A: Transverse section stained with a cresyl violet acetate solution. B: Schematic diagram showing the location of descending motor tracts [23]: the corticospinal tract (CST; green), rubrospinal tract (RbST; red), and reticulospinal tract (RtST; blue).

### Spike Frequency in HLA During ADS

The profiles of spikes recorded in HLA and used to trigger stimulation during each hour of ADS remained consistent over the 3-hours of ADS (Figure 5, Left). Additionally, there were no significant changes in the HLA spike frequency in any group after 3 hours of ADS relative to the 1st hour of ADS (Figure 5, Right). While there were minor within-group changes in average spike frequency during the second and third hour of ADS in some groups relative to the first hour of ADS, these changes were not significant.

**Figure 5.**
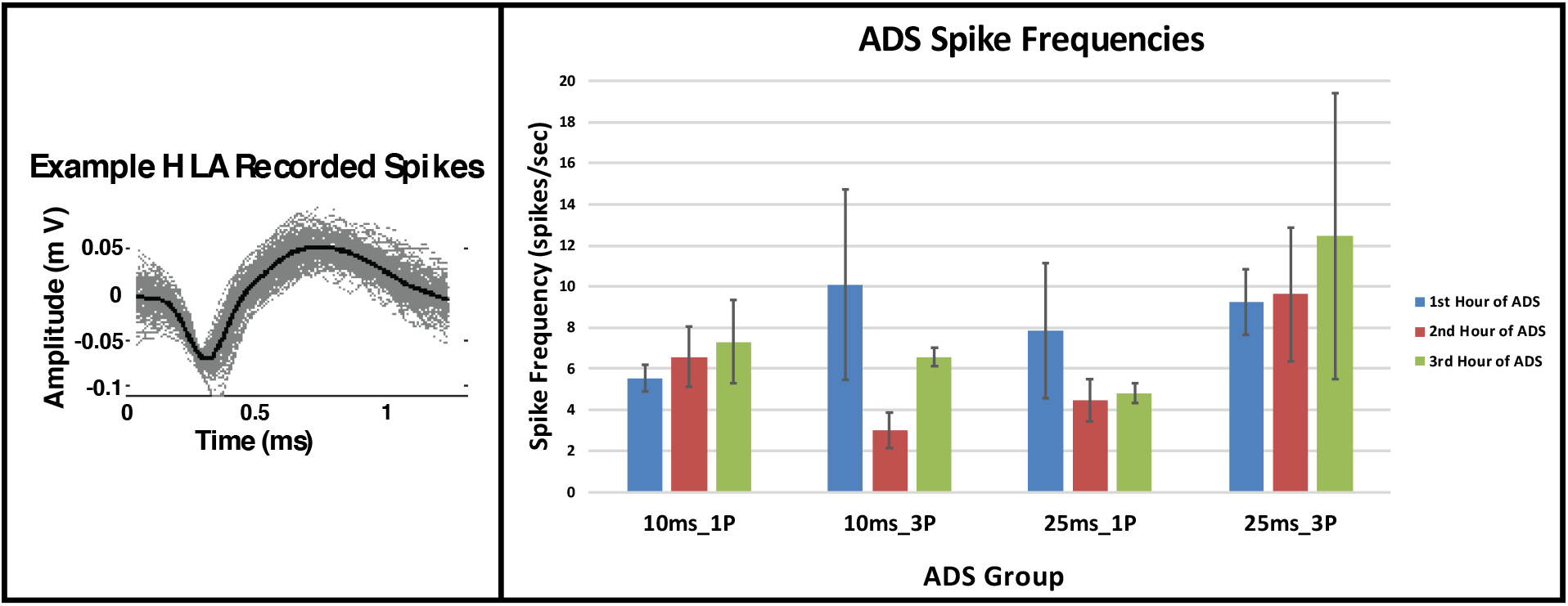
Extracellular spikes recorded in hindlimb motor cortex (HLA) during the ADS sessions. **Left:** Exemplar spike profiles recorded from layer V of hindlimb motor cortex from one rat. These spikes were used to trigger ISMS pulses in the ventral horn of the lumbar spinal cord during ADS. Grey spikes are all spikes of similar spiking profile recorded from layer V and the solid black spike is the average of these recorded spikes. **Right:** Average spike frequencies of recorded spikes for each parameter group during each hour of ADS. Data were derived from 5-minute recordings at the end of each hour of ADS.

### ISMS and ICMS Current Intensity

The average ISMS movement thresholds and ADS stimulus intensities for each group are shown in Table II. The average ISMS stimulus intensity required to evoke a hindlimb movement (movement threshold) during spinal cord mapping was 9.00 ± 5.90 μA. The average ISMS intensity used for ADS bouts was 4.50 ± 2.95 μA (i.e., ADS intensity). There was no statistical difference in either ISMS movement threshold intensities nor ISMS intensity used during ADS bouts between groups (Table 2; *p* > 0.05). Since movement could not be evoked during ICMS mapping in SCI rats, a fixed intensity of 100 μA was used for ICMS during the 5-minute baseline and test bouts.

**Table II.** Stimulation Movement Thresholds and Intensities Used. Mean and standard deviations at four weeks post-SCI during the ADS procedure. ISMS threshold was recorded during ISMS mapping at movement threshold. ADS intensity was 50% of ISMS threshold. ICMS intensity was set at 100 μA and used during the 5-minute baseline and test bouts.

| Group | ISMS Threshold (μA) | ADS Intensity (μA) | ICMS Intensity (μA) |
| --- | --- | --- | --- |
| 10ms_1P | 7.14 ± 2.27 | 3.57 ± 1.13 | 100.00 ± 0.00 |
| 25ms_1P | 8.86 ± 5.98 | 4.43 ± 2.99 | 100.00 ± 0.00 |
| 10ms_3P | 10.80 ± 8.79 | 5.40 ± 4.39 | 100.00 ± 0.00 |
| 25ms_3P | 10.00 ± 7.07 | 5.00 ± 3.54 | 100.00 ± 0.00 |
| Combined | 9.00 ± 5.90 | 4.50 ± 2.95 | 100.00 ± 0.00 |

### EMG Activity

ICMS-evoked EMG activity was never observed before or after ADS (i.e., during baseline and test bouts, no EMG potentials were greater than two standard deviations above the baseline EMG activity). During ADS, ongoing EMG amplitude fluctuated periodically, but no consistent changes were evident. The following qualitative observations (and their incidence) are of note: 1) EMG activity was present in each hindlimb muscle and EMG amplitude increased over the 3 hours of ADS (n = 6 instances), 2) EMG activity was present during the 1^st^ hour of ADS but was not present after the 2^nd^ or 3^rd^ hour of ADS (n = 3 instances), 3) EMG activity was never present over the 3 hours of ADS (n = 10 instances), and 4) EMG activity increased during the 2^nd^ hour of ADS but was not present after the 3^rd^ hour of ADS (n = 1 instance). These observations did not occur systematically and thus the data were not analyzed further.

### Post-stimulus spike histograms

The ICMS-evoked spiking activity was recorded for each channel and analyzed over four dorsoventral sectors delineated by depth below the surface of the spinal cord. Exemplar ICMS-evoked spikes are displayed in Figure 6. After 3 hours of ADS, post-stimulus spike histograms from the 10ms_1P group (Figure 7) showed qualitatively that ICMS-evoked spiking activity increased at a latency of ~10 ms from onset of ICMS (i.e. 0 ms), especially within the ventral sector.

**Figure 6.**
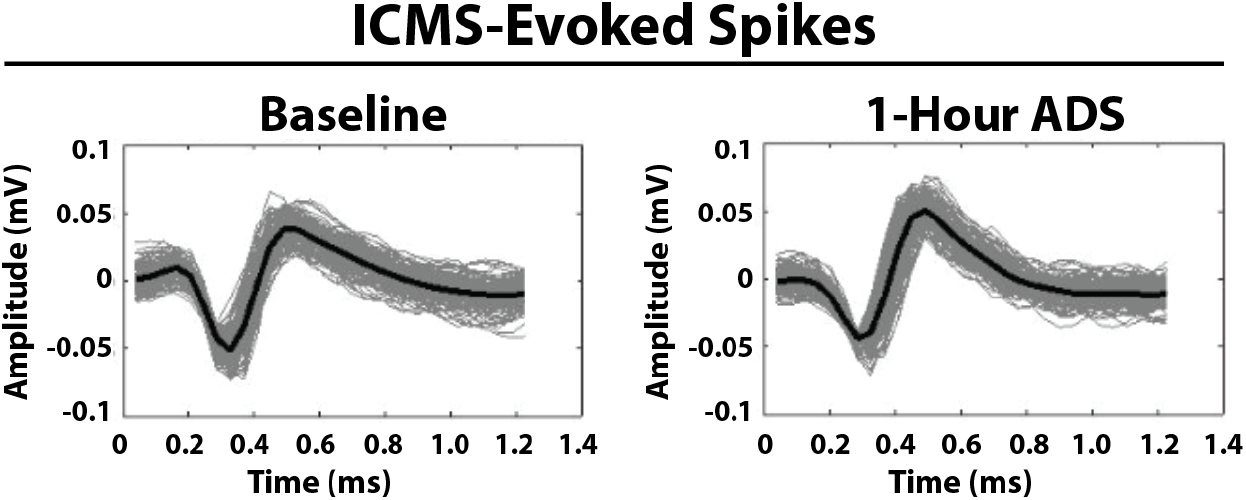
Exemplar ICMS-evoked spikes. Recorded ICMS-evoked spikes from the Lower Intermediate Sector of the hindlimb spinal cord from one rat in the 25ms_1P group before (Baseline) and after 1-Hour of ADS. The solid black spike is the average of these recorded spikes. Spike profiles remained similar between bouts of ADS therapy.

**Figure 7.**
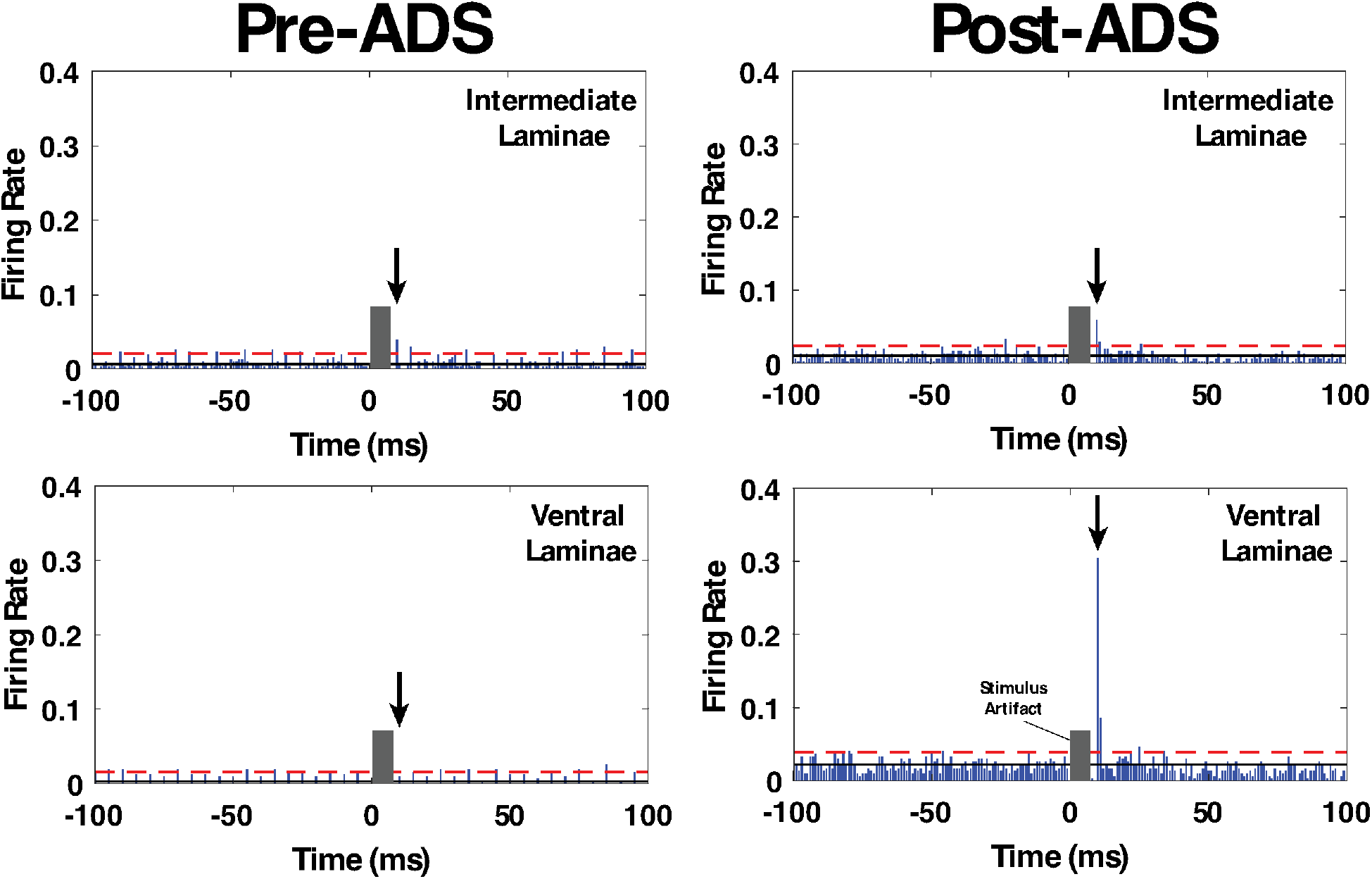
Exemplar post-stimulus spike histograms from rats in the 10ms_1P group before and after 3 hours of ADS. Post-ICMS spikes occurred during a short latency period from 10-12 ms, and a long-latency period from 24-36 ms. Black arrow indicates the short-latency bin with the highest ICMS-evoked FR, and thus used for pre/post ADS comparison. The number of recorded ICMS-evoked spikes during the short-latency period increased in the ventral laminae (lower right) after 3 hours of ADS. Solid black line is average FR before onset ICMS (i.e., pre-ICMS FR). Red dashed line is 2x standard deviations above pre-ICMS FR.

### ICMS-Evoked Maximum Firing Rate at Short Latency

Within each group, maximum firing rate was examined at the end of each of the one-hour ADS sessions for each of the four dorsoventral quadrants. Maximum FR during the short-latency period typically occurred at 10 ms (all test bouts; mean ± SE = 10.34 ± 0.07 ms; median = 10.00 ms) relative to ICMS, but occasionally at 11 and 12 ms.

In each of the four groups, FR increased in one or more dorsoventral sectors of the cord after ADS compared with baseline FR (Figure 8). When significant increases in FR occurred, the FR usually remained significantly increased from baseline. For example, if an increase in FR occurred 2 hours after ADS, FR usually remained significantly increased after 3 hours of ADS.

**Figure 8.**
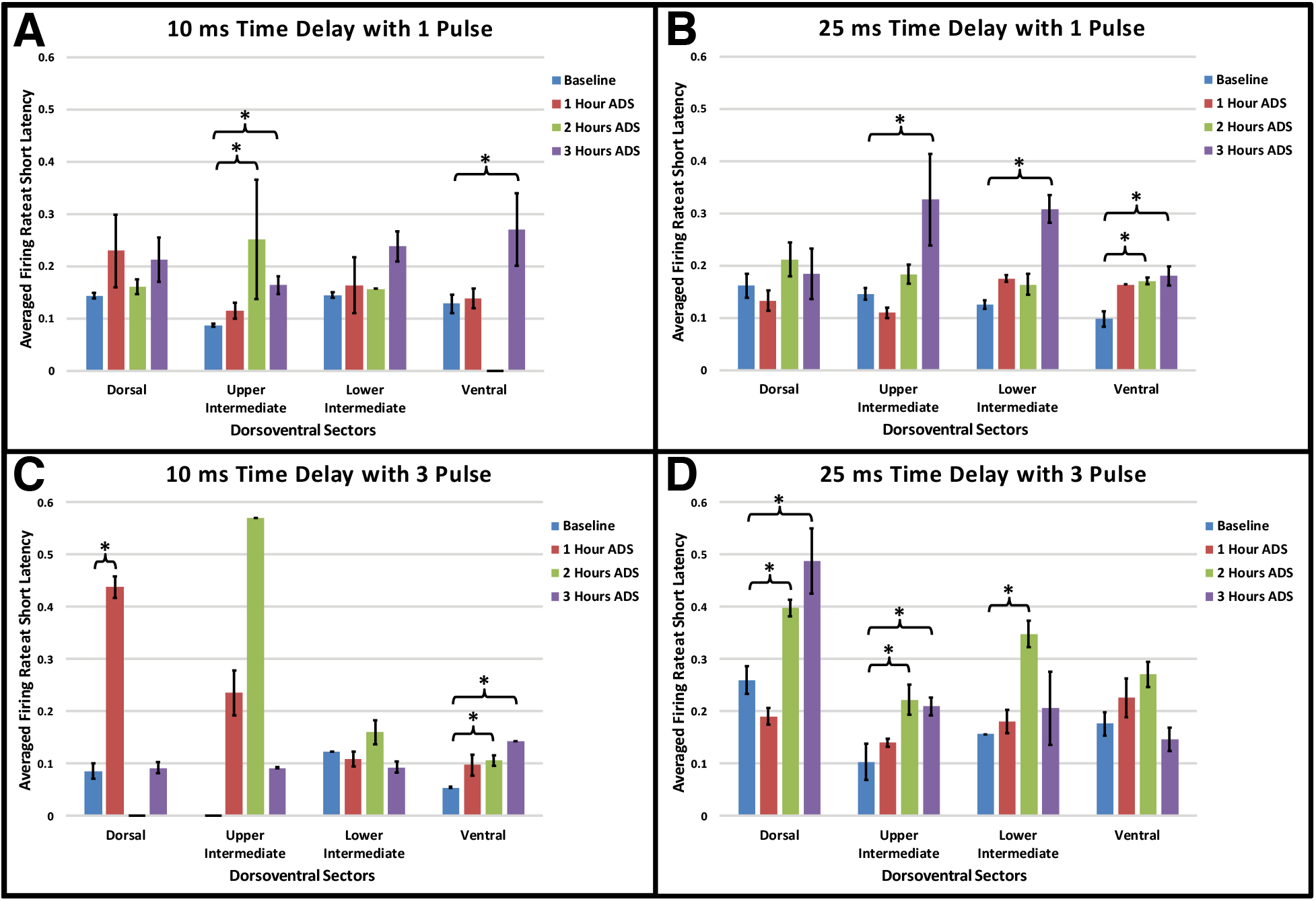
Maximum ICMS-evoked firing rates (i.e., means + std err) during the short-latency period for each dorsoventral sector at baseline and after each hour of ADS. Generally, the 10 ms post-ICMS bin was used for this analysis. **A)** 10ms_1P ADS group, **B)** 25ms_1P ADS group, **C)** 10ms_3P ADS group, and **D)** 25ms_3P ADS group. * = Significant increase in maximum firing rate when compared to baseline recordings (*p* < 0.05).

While systematic differences between groups with different spike-stimulus latencies or number of pulses was not observed, some trends are worth noting. With a 10ms time delay protocol, increased short latency FR was more prevalent in the ventral horn; with a 25ms delay, FR increases were less specific. Also, while a single pulse protocol most often resulted in FR increases in the upper intermediate and ventral sectors, a 3-pulse protocol resulted in ventral FR increases with a short delay and more dorsal FR increases with the longer delay.

### ICMS-Evoked Maximum Firing Rate at Long Latency

Within each group, maximum firing rate was examined at the end of each of the one-hour ADS sessions for each of the four dorsoventral quadrants. Maximum FR during the long-latency period occurred at about 25 ms (all test bouts; mean ± SE = 24.88 ± 0.65 ms; median = 25.00 ms).

As observed in the short latency period, in each of the four groups FR increased in one or more dorsoventral sectors of the cord after ADS compared with baseline FR (Figure 9). Again, when significant increases in FR occurred, the FR usually remained significantly increased from baseline. However, increases in long latency FR occurred more commonly after 2 hours of ADS and as early as 1 hour of ADS.

**Figure 9.**
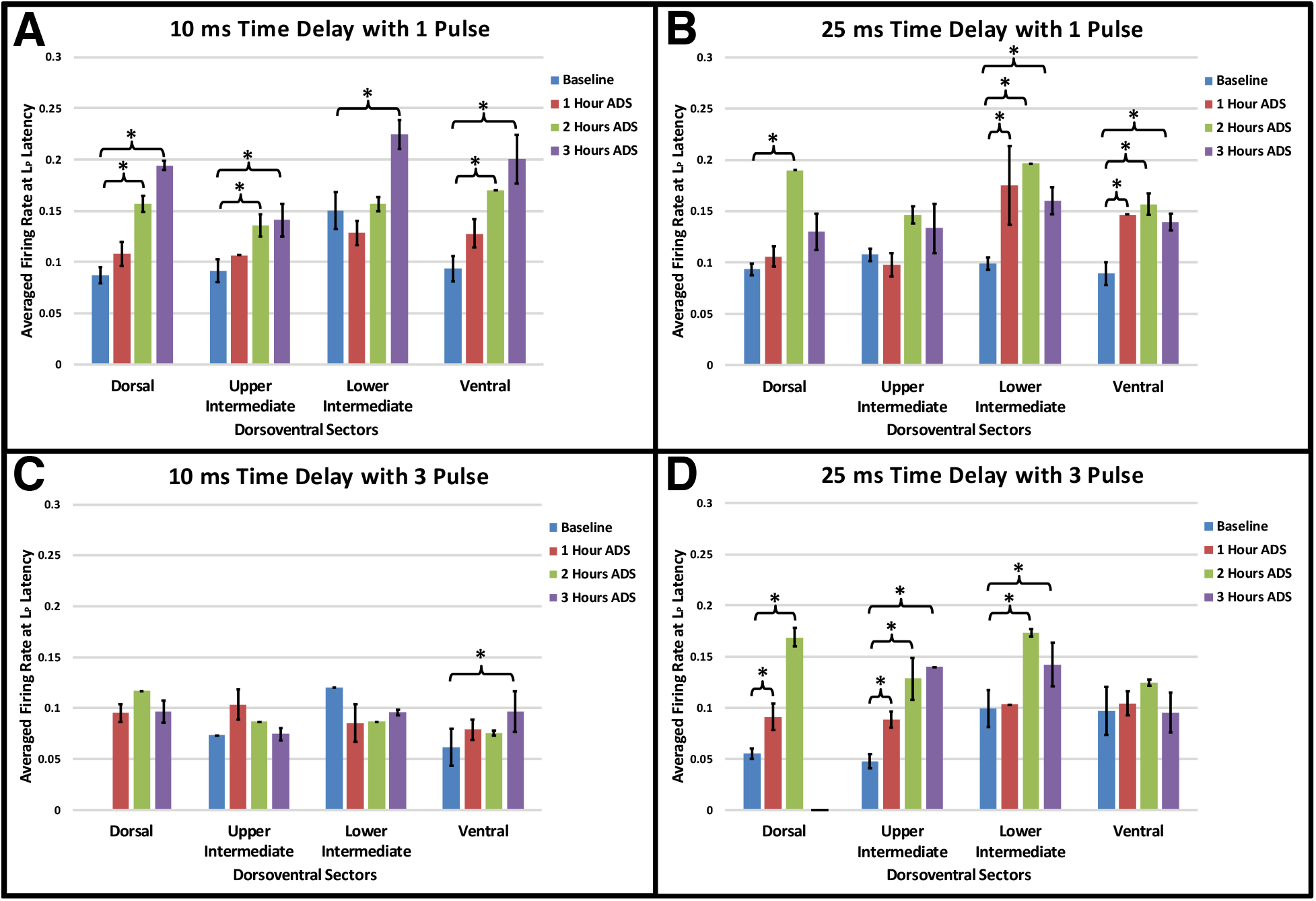
Maximum ICMS-evoked firing rates (i.e., means + std err) during the long-latency period for each dorsoventral sectors at baseline and after each hour of ADS. Generally, the 24-36 ms post-ICMS bins was used for this analysis. **A)** 10ms_1P ADS group, **B)** 25ms_1P ADS group, **C)** 10ms_3P ADS group, and **D)** 25ms_3P ADS group. * = Significant increase of firing rate when compared to baseline recordings (*p* < 0.05).

In general, with a 10 ms delay protocol, long latency FR increased in all sectors using a single pulse, but only increased in the ventral sector using a 3-pulse stimulus. With a 25 ms delay protocol FR increased predominantly in the more ventral sectors using a single pulse and increased in more dorsal sectors with three pulses.

## Discussion

The primary goal of this study was to determine whether activity dependent stimulation (ADS) using different spike-stimulus delays and number of ISMS pulses alters synaptic efficacy in remaining intact cortical descending motor pathways in an anesthetized rat model of SCI. By pairing sites of similar functional topography in the hindlimb motor cortex and hindlimb spinal cord and synchronizing the delivery of ISMS in the ventral horn with spontaneous neural activity in motor cortex, the synaptic efficacy of descending motor pathways increased, as evidenced by an increase in cortically-evoked spikes in spinal cord neurons at specific post-ICMS latencies. For the first time, cortically-evoked spikes were recorded from multiple dorsoventral sectors in an acute ADS paradigm. Importantly, efficacy in descending motor pathways was increased throughout four dorsoventral sectors of the hindlimb spinal cord, depending on the combination of time delay and pulse number used. Any increase in firing rate (FR) in the Ventral sector is considered important for functional recovery since it may indicate increased synaptic efficacy in pathways that directly influence motor neurons. The observed increase of FR at short latencies in the Ventral sector was significant after several post-ADS test bouts whether a 10 ms or 25 ms delay was used. Here we discuss the likely motor pathways affected by ADS and the functional impact these synaptic efficacy changes could have in motor control.

### Spike-Stimulus Delays during ADS are Consistent with STDP-Based Strengthening

In spike timing-dependent plasticity (STDP), the timing between recorded action potentials and triggered stimulation (i.e., ADS) has been shown to determine the polarity and magnitude (i.e., potentiation or depression) of any change in post-synaptic potentials [5, 24, 25]. McPherson *et al.* demonstrated that ADS can induce neural plasticity, with a time delay of ~10 ms, that improves behavioral recovery after SCI by synchronizing ISMS below the injury with the arrival of functionally related volitional motor commands signaled by muscle activity in the impaired forelimb [1]. The time delays used during ADS in this study fell within the range of neurophysiologically relevant latencies (i.e., ~25 ms or less), where robust enhancement of synaptic strength was determined *in vitro* [26] and *in vivo* [7] studies.

Although a 3-pulse ISMS train (typical protocol for activity-triggered ISMS) is problematic for a simple model of spike-timing relationships, we still observed significant increases in FR when the pulse train spanned approximately 10 – 20 ms after the cortical spike. With long delays and multiple pulses, the synaptic conditioning effects may be compromised since the later pulses fall outside of the optimal window for long term potentiation.

### Short-Latency Neuronal Spikes after Acute ADS

We showed in a previous study that short-latency (10-12 ms) spikes were still evoked from cortex after a moderate spinal cord contusion identical to that used here [16]. These post-injury spikes were most likely evoked via cortico-reticulospinal fibers, based on the fast conduction velocities measured and the observation that the ventrolateral funiculus, where reticulospinal fibers descend, was spared. It is likely that these fibers contribute to the increase in short latency cortically evoked activity after application of ADS.

*Reticulospinal Tract:* The RtSt is strongly involved in locomotor coordination in healthy animals [27] and plays an important role in locomotion after SCI [28]. Spared reticulospinal fibers are known to sprout below an experimental SCI [29]. The reticulospinal tract terminates mainly in the grey matter of lamina VII and VIII in the hindlimb spinal cord of the rat [27, 30], which corresponds to the Lower Intermediate sector and part of the Ventral sector of the spinal cord described in the present study. Thus, it is likely that the RtSt, activated via the spiking neurons in cortex, participated in the ADS-induced increase in FR.

*Propriospinal Fibers:* The increase in short-latency FR in the Ventral sector (and occasionally in the Lower Intermediate sector) after ADS could be mediated by propriospinal fibers activated by the reticulospinal tract fibers that terminate onto long descending propriospinal neurons involved in the coordination between forelimb-hindlimb and left-right limbs [31]. Polysynaptic connections onto long descending propriospinal pathways [32] could have been activated in chorus with cortico-reticulospinal pathways at the Short latency period. The increased FR at short-latency seen in the Dorsal and Upper Intermediate sectors after ADS may suggest a strengthening of the cortico-reticulospinal synapses onto these intact long descending propriospinal pathways. Any increase in FR in the Dorsal and Upper Intermediate sectors may indicate increased synaptic efficacy in interneurons that influence the overall neural circuit that influences motor neurons.

### Long-Latency Neuronal Spikes & EMG Activity after Acute ADS

In a previous study, it was shown that the ICMS-evoked long-latency spinal cord spike activity in a healthy, uninjured rat was represented in a broad distribution beginning ~20 ms after ICMS onset, then overlapping with the onset of EMG activity at ~25 ms, and peaking around ~30 ms (median of peak long-latency response, or L_p_) [16]. After injury, this spiking pattern was disrupted, as the FR at long latencies was reduced substantially, and EMG activity could not be evoked. While FR occasionally rose above baseline, significant spiking was restricted to only one or two millisecond bins within this broad time period. This result was replicated after SCI in the present study, as the FR of long-latency neuronal spikes rose above baseline for only a few time bins within the broad range of 24-36 ms. It was argued that the evoked spikes with long latencies were most likely from neurons activated by the corticospinal and (cortico-) rubrospinal tracts due to the known conduction velocities of these tracts.

The corticospinal tract (CST) terminates mainly in the Dorsal and Upper Intermediate sectors in the hindlimb spinal cord of the rat, with a few fibers terminating in all dorsoventral sectors [27, 33, 34]. Direct corticomotoneuronal connections are unsupported in rats [23, 35]. The rubrospinal tract (RbST) primarily terminates in the intermediate grey matter corresponding to Upper and Lower Intermediate sectors [36]. The influence of CST and RbST on spiking in the Ventral sector is likely exerted via interneurons in dorsal and intermediate laminae. In the present study, it is assumed that the CST, descending primarily in the dorsomedial funiculus in the rat, was heavily damaged after the SCI. Thus, any ICMS-driven spikes in the Ventral sector at baseline, and any increases in FR after ADS bouts could have been mediated by polysynaptic activation via spared fiber tracts, including RbST, dorsolateral CST, ventral CST, reticulospinal tract, and vestibulospinal tract.

### Impact of Acute Plasticity on Recovery

The increased FR of spinal cord neurons in some conditions as early as 1 hour after ADS is a novel and significant finding that may have clinical relevance. These results may drive future electrical stimulation-based therapies. Currently, most electrical stimulation-based therapies are contingent on the maintained delivery of stimulation. The results seen here show that enhancement of descending motor pathways can occur after an acute application of ADS therapy. Future studies will have to determine if enhanced synaptic efficacy is maintained post-therapy, the functional significance of this plasticity, its effect on functional motor recovery in the chronic condition, and the potential for behavioral improvement compared to that found after less-invasive electrical stimulation-based therapies.

### Limitations of Study

It is important to note the limitations in the present study. The main technical limitation was the necessity to remove and reinsert the microelectrodes between the 1-hour ADS sessions and the 5-minute test bouts. When removing and reinserting the electrodes, there was a possibility that the microelectrodes were not reinserted into the exact position as before. In addition, there could be neuronal damage with each reinsertion of the electrode. The ICMS and ISMS mapping did reduce the amount of variation in site location between each ADS session and test bout. The use of two micromanipulators (one for each electrode) also aided with the consistency of electrode insertion and placement on a micron scale. The consistency in recorded HLA spiking profiles over the ADS sessions also indicates that the recording microelectrode inserted in HLA was consistently in the same site each time. A reliable switchable headstage that can automatically switch between recording and stimulation would allow for the microelectrodes to remain in their original sites. These were not available at the time of this study.

There was an absence or decrease in spiking activity at various time points as seen in Figures 8A, 8C, and 9D. This decrease in FR could be a physiological significance such as suppression; however, our assumption is that the spike-to-noise ratio was so low that no spikes could be registered via our current spike sorting methods. This would be caused by any significant noise in the signal during recording.

Similarly, there was a difference in baseline FR between groups at some dorsoventral sectors. This could indicate that the spinal neuronal population recorded and analyzed was undersampled. For this reason, between-group comparisons were not conducted. However, within-group comparisons were conducted since similar neuronal populations could be compared, which increases the likelihood of detecting significant differences in FR over time. Future experiments will strive to better identify the sub-population of neurons contributing to spikes recorded in each dorsoventral sector. In addition, increasing the number of subjects and number of test bouts will help reduce the variability of recorded spikes between-groups. As this study included all evoked spikes from any intact descending pathways, future studies are planned to examine the contribution of specific pathways. This too will reduce any variability in the sub-populations recorded from different dorsoventral sectors.

It is possible that different ICMS parameters during the test bouts may be more effective for assessing EMG activity and evoked hindlimb movement after ADS therapy. Trains consisting of 13 stimulation pulses are most commonly used for evoking movement during motor cortex mapping procedures [13, 14] and only 3 stimulation pulses were used during ICMS test bouts in this study. As a result, the shorter ICMS pulse train may not have allowed adequate temporal summation to activate the motor neurons (i.e., reach motor movement threshold) in the hindlimb spinal cord. Additionally, a more chronic application of ADS may be needed for the plasticity to result in functional motor recovery.

Finally, the present results were based on the topographic location of the electrodes through the dorsoventral depths of the spinal cord grey matter. The extracellular spike recordings were derived from a heterogeneous group of spinal cord neurons that were not specifically characterized. Likewise, the evoked spikes were the result of several polysynaptic pathways from hindlimb motor cortex, including ventral corticospinal, rubrospinal, reticulospinal and vestibulospinal tracts. The contribution of each of these spared pathways on the ability to drive post-SCI plasticity with ADS will need to be addressed, potentially using selective tract lesions in a similar neurophysiological preparation.

## Conclusion

The significant increase in cortically-evoked firing of spinal cord neurons after acute ADS indicates a strengthening of the remaining descending pathways. ADS resulted in greater responsiveness in neurons throughout the dorsoventral depths of the spinal cord, and under a wide range of timing parameters. These results may suggest appropriate parameters and locations for chronic administration of cortically-driven ADS that can be tested with direct measurement of behavioral outcomes. This may provide further guidance for future neurorehabilitation and neuromodulatory interventions that may better contribute to enhanced recovery of motor function after moderate spinal cord injuries.

## Data Availability Statement

Data available on request from the authors.

## Conflict of Interest Statement

The authors declare no competing financial interests.

## Acknowledgments & Funding Statement

This work was supported by the Paralyzed Veterans of America Research Foundation #3068, The Ronald D. Deffenbaugh Family Foundation, NIH/NINDS R01 NS030853, T32 Neurological and Rehabilitation Sciences Training Program, and NIH/NINDS F31 NS105442.

## Notes

### Competing Interest Statement

The authors have declared no competing interest.

### Summary of Updates

Introduction, Results and Discussion has been updated.

## References

1. McPherson, J.G., R.R. Miller, and S.I. Perlmutter, Targeted, activity-dependent spinal stimulation produces long-lasting motor recovery in chronic cervical spinal cord injury. Proc Natl Acad Sci U S A, 2015. 112(39): p. 12193–8.

2. Gad, P., et al., Forelimb EMG-based trigger to control an electronic spinal bridge to enable hindlimb stepping after a complete spinal cord lesion in rats. J Neuroeng Rehabil, 2012. 9(38): p. 38.

3. Wenger, N., et al., Spatiotemporal neuromodulation therapies engaging muscle synergies improve motor control after spinal cord injury. Nat Med, 2016. 22(2): p. 138–45.

4. Capogrosso, M., et al., A brain-spine interface alleviating gait deficits after spinal cord injury in primates. Nature, 2016. 539(7628): p. 284–288.

5. Bi, G. and M. Poo, Synaptic Modifications in Culutured Hippocampal Neurons: Dependence on Spike Timing, Synaptic Strength, and Postsynaptic Cell Type. The Journal of Neuroscience, 1998. 18(24): p. 10464–10472.

6. Jackson, A., J. Mavoori, and E.E. Fetz, Long-term motor cortex plasticity induced by an electronic neural implant. Nature, 2006. 444(7115): p. 56–60.

7. Nishimura, Y., et al., Spike-timing-dependent plasticity in primate corticospinal connections induced during free behavior. Neuron, 2013. 80(5): p. 1301–9.

8. Behrmann, D.L., et al., Spinal Cord Injury Produced by Consistent Mechanical Displacement of the Cord in Rats: Behavioral and Histological Analysis. Journal Of Neurotrauma, 1992. 9(3): p. 197–217.

9. Basso, D.M., M.S. Beattie, and J.C. Bresnahan, A sensitive and reliable locomotor rating scale for open field testing in rats. J Neurotrauma, 1995. 12(1): p. 1–21.

10. Scheff, S.W., et al., Experimental modeling of spinal cord injury: characterization of a force-defined injury device. J Neurotrauma, 2003. 20(2): p. 179–93.

11. Krizsan-Agbas, D., et al., Gait Analysis at Multiple Speeds Reveals Differential Functional and Structural Outcomes in Response to Graded Spinal Cord Injury. Journal of Neurotrauma, 2014. 31(9): p. 846–856.

12. Borrell, J.A., et al., A 3D map of the hindlimb motor representation in the lumbar spinal cord in Sprague Dawley rats. J Neural Eng, 2017. 14(1): p. 016007.

13. Frost, S.B., et al., Reliability in the location of hindlimb motor representations in Fischer-344 rats: laboratory investigation. J Neurosurg Spine, 2013. 19(2): p. 248–55.

14. Frost, S.B., et al., Output Properties of the Cortical Hindlimb Motor Area in Spinal Cord-Injured Rats. J Neurotrauma, 2015. 32(21): p. 1666–73.

15. Lewicki, M.S., A review of methods for spike sorting: the detection and classification of neural action potentials. Network, 1998. 9(4): p. R53–78.

16. Borrell, J.A., et al., Effects of a contusive spinal cord injury on cortically-evoked spinal spiking activity in rats. J Neural Eng, 2020. 17.

17. Bi, G. and M. Poo, Synaptic modification by correlated activity: Hebb’s postulate revisited. Annu Rev Neurosci, 2001. 24: p. 139–66.

18. Moritz, C.T., et al., Forelimb movements and muscle responses evoked by microstimulation of cervical spinal cord in sedated monkeys. J Neurophysiol, 2007. 97(1): p. 110–20.

19. Gilerovich, E.G., et al., Morphofunctional Characteristics of the Lumbar Enlargement of the Spinal Cords in Rats. Neuroscience and Behavioral Physiology, 2008. 38(8): p. 855–860.

20. Molander, C., Q. Xu, and G. Grant, The cytoarchitectonic organization of the spinal cord in the rat. I. The lower thoracic and lumbosacral cord. J Comp Neurol, 1984. 230(1): p. 133–41.

21. Hudson, H.M., et al., Properties of primary motor cortex output to hindlimb muscles in the macaque monkey. J Neurophysiol, 2015. 113(3): p. 937–49.

22. Averna, A., et al., Differential Effects of Open- and Closed-Loop Intracortical Microstimulation on Firing Patterns of Neurons in Distant Cortical Areas. Cereb Cortex, 2020. 30(5): p. 2879–2896.

23. Lemon, R.N., Descending pathways in motor control. Annu Rev Neurosci, 2008. 31: p. 195–218.

24. Markram, H., et al., Regulation of Synaptic Efficacy by Coinicidence of Postsynaptic APs and EPSPs. Science, 1997. 275: p. 213–215.

25. Fung, T.K., C.S. Law, and L.S. Leung, Associative spike timing-dependent potentiation of the basal dendritic excitatory synapses in the hippocampus in vivo. J Neurophysiol, 2016. 115(6): p. 3264–74.

26. Feldman, D.E., The spike-timing dependence of plasticity. Neuron, 2012. 75(4): p. 556–71.

27. Mitchell, E.J., et al., Corticospinal and Reticulospinal Contacts on Cervical Commissural and Long Descending Propriospinal Neurons in the Adult Rat Spinal Cord; Evidence for Powerful Reticulospinal Connections. PLoS One, 2016. 11(3): p. e0152094.

28. Schucht, P., et al., Anatomical Correlates of Locomotor Recovery Following Dorsal and Ventral Lesions of the Rat Spinal Cord. Experimental Neurology, 2002. 176(1): p. 143–153.

29. Ballermann, M. and K. Fouad, Spontaneous locomotor recovery in spinal cord injured rats is accompanied by anatomical plasticity of reticulospinal fibers. Eur J Neurosci, 2006. 23(8): p. 1988–96.

30. Matsuyama, K., et al., Morphology of Single Pontine Reticulospinal Axons in the Lumbar enlargement of the Cat: A Stud Using the Anterograde Tracer PHA-L. The Journal of Comparative Neurology, 1999. 410: p. 413–430.

31. Frigon, A., The neural control of interlimb coordination during mammalian locomotion. J Neurophysiol, 2017. 117(6): p. 2224–2241.

32. Skinner, R.D., et al., Cells of origin of long descending propriospinal fibers connecting the spinal enlargements in cat and monkey determined by horseradish peroxidase and electrophysiological techniques. J Comp Neurol, 1979. 188(3): p. 443–54.

33. Casale, E.J., A.R. Light, and A. Rustioni, Direct projection of the corticospinal tract to the superficial laminae of the spinal cord in the rat. J Comp Neurol, 1988. 278(2): p. 275–86.

34. Akintunde, A. and D.F. Buxton, Differential sites of origin and collateralization of corticospinal neurons in the rat: a multiple fluorescent retrograde tracer study. Brain Res, 1992. 575(1): p. 86–92.

35. Yang, H.W. and R.N. Lemon, An electron microscopic examination of the corticospinal projection to the cervical spinal cord in the rat: lack of evidence for cortico-motoneuronal synapses. Exp Brain Res, 2003. 149(4): p. 458–69.

36. Brown, L.T., Rubrospinal projections in the rat. J Comp Neurol, 1974. 154(2): p. 169–87.

